# Sex and gender differences in perivascular space in early adolescence

**DOI:** 10.1101/2025.09.30.679416

**Authors:** Carinna Torgerson, Hedyeh Ahmadi, Haoyu Lan, Jessica Morrel, Megan Herting, Jeiran Choupan

## Abstract

Perivascular spaces (PVS) surrounding cerebral blood vessels play an important role in the blood-brain barrier and glymphatic system. Although it was once thought that PVS were either absent or too small to be seen or quantified during healthy development with MRI, recent studies have found visible, quantifiable PVS exist throughout the white matter of the cerebrum in childhood and adolescence. As a result, researchers have begun to explore individual differences, including potential sex-based variations in developing PVS. Meta-analyses in adults have shown that PVS are larger on average in males than in females, and several studies have shown a similar relationship in children. In contrast, no studies to date have examined the association between gender and PVS at any age. This cross-sectional study examined 6,538 youths from a large, nationwide sample of 9- to 11-year-olds in the U.S. to examine the relationship between sex, felt-gender, and PVS count and volume. Using a model-building approach, we conducted a series of linear mixed-effects models to determine the maximum variance explained in PVS count and volume, including age, pubertal development status, race, parent education, BMI z-score, and regional white matter volume, while also adjusting for MRI scanner and site. BMI z-score, age, and parent education were significant predictors of both PVS volume and count. Adding sex to the model improved model fit in all regions, and the further addition of felt-gender significantly improved model fit for PVS count in 5/6 regions of interest. Moreover, we found increases in PVS volume and count were associated with reduced executive function, learning, and memory. As the first study to report an association between felt-gender and PVS, our findings demonstrate the importance of considering gender in addition to sex as a potential source of structural variance in PVS in adolescents.

## Introduction

Perivascular spaces (PVS) are extracellular spaces surrounding blood vessels which allow transport-mediated exchange of solutes between the blood and cerebrospinal fluid (CSF) (1). MRI-visible PVS were long considered absent or rare in children (1). Most studies of PVS in children therefore focus on specific patient populations (2,3) or assess lifespan development of PVS using both children and adults (4,5). Nonetheless, a few recent studies have shown that PVS are common and measurable in healthy adolescents (ages 12-21) and that significant PVS development occurs during adolescence (6,7). Two studies have also reported sex differences in adolescents, with PVS occupying a larger proportion of regional brain volume on average in boys compared to girls (4,7), which aligns with studies in adults showing larger PVS in males (4,8,9). Beyond sex and age, PVS structure has been associated with several cardiovascular health factors, including body mass index (BMI) (8,10) Although no studies to date have examined the relationship between PVS and gender, transgender people and gender minorities experience disproportionate risk for poor vascular health and also experience excess cardiovascular morbidity not fully explained by these individual risk factors (11). In adults, enlarged PVS in the centrum semiovale are associated with worse general intelligence, executive function, language, and memory (12); however, the literature in adults typically describes enlarged PVS as a result of neurodegeneration or pathology (13,14). It is unclear whether PVS are also associated with cognition in neurologically healthy adolescents. Therefore, this study aimed to explore the demographic determinants of PVS and the impact of PVS structure on cognition in early adolescents ages 9-11 years-old. We hypothesized that we would be able to detect and quantify PVS in all regions for all participants, that we would find a larger average number of PVS (e.g., count) in male youth compared to female youth, and that sex would explain more variance in PVS than felt-gender.

## Methods

### Participants

The current study uses data from the ongoing Adolescent Brain Cognitive Development (ABCD) Study^®^, which includes 11,868 children at 21 sites around the United States (15,16). More information on the inclusion criteria and data used is available in the Supplemental Methods. Data was obtained from the MR images and demographic questionnaires collected in the first two years of the ABCD Study. In accordance with the study design, imaging data was collected at the baseline visit (ages 9–10 years), whereas gender questionnaire responses were collected at the first follow-up visit (ages 10–11 years) (15,17). We then restricted our sample to participants at the 21 primary study sites who had an X/Y allele frequency ratio matching their reported sex, met the ABCD consortium quality control standards for raw imaging or FreeSurfer reconstruction, had no incidental findings warranting clinical referral after inspection by a radiologist, and had no missing data (Figure 1). Since some ABCD Study sites targeted enrollment of twins, triplets, and siblings, we restricted our sample to one child per family to reduce familial bias. As previously published (18), to improve our ability to detect associations between felt-gender and PVS, in cases where siblings’ scores differed we selected the child with the lowest felt-gender score. If siblings had the same felt-gender score, one child was chosen randomly. After applying these inclusion and exclusion criteria, our sample consisted of 6,538 children (3,228 female). A complete description of the final sample for the current study can be found in Table 1.

**Figure 1.**
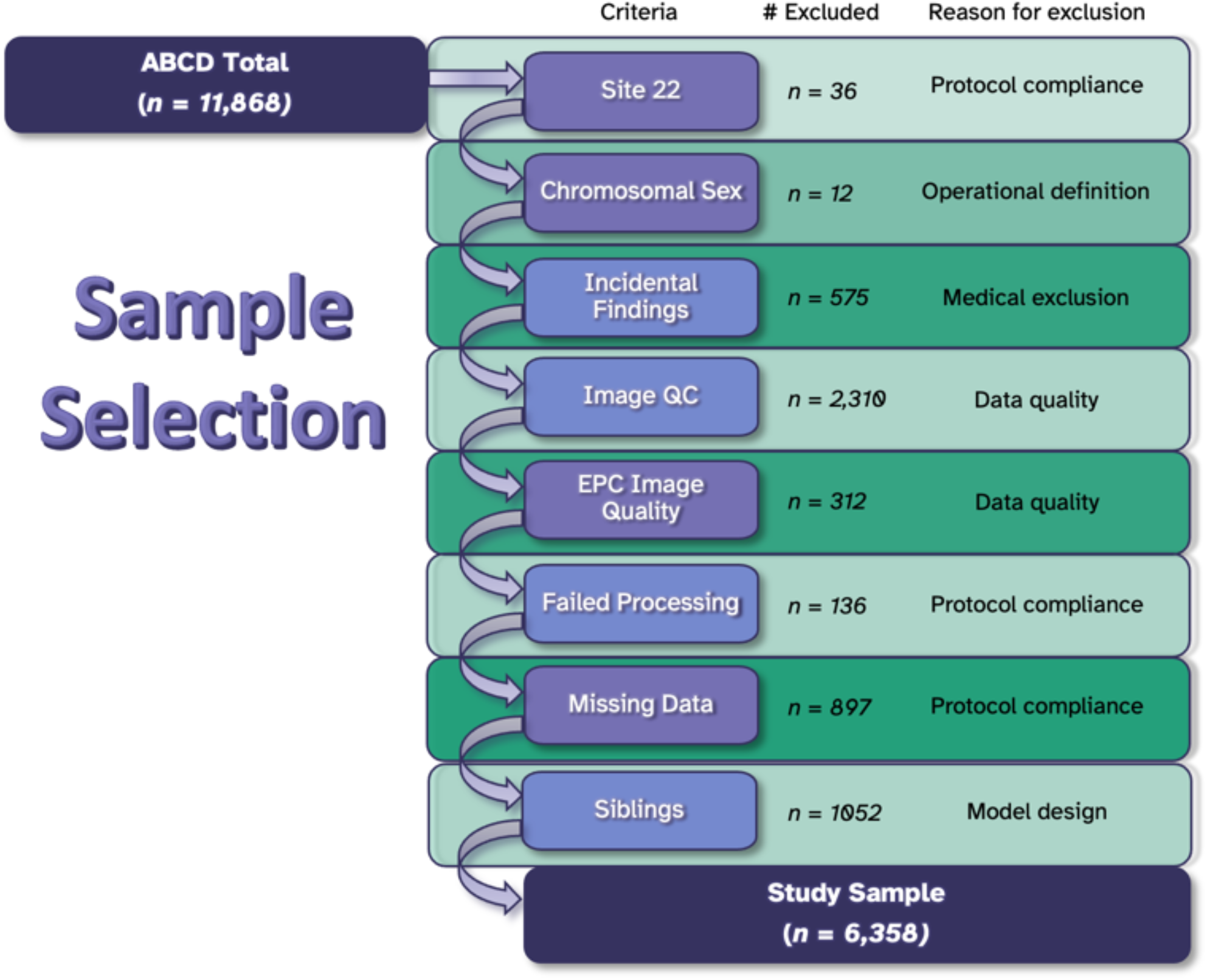
Flow diagram demonstrating the sample selection process along with number of participants excluded and the reasons for exclusion.

**Table 1.** Comparison of the demographics of the overall ABCD sample with the study sample.

|  | ABCD<br>(N=11868) | Study Sample<br>(N=6538) |
| --- | --- | --- |
| **Sex** |  |  |
| Female | 5677 (47.8%) | 3228 (49.4%) |
| Intersex-Male | 3 (0.0%) | 0 (0%) |
| Male | 6188 (52.1%) | 3310 (50.6%) |
| **<sup>a</sup> Felt-Gender** |  |  |
| Mean (SD) | 4.80 (0.509) | 4.78 (0.534) |
| Median [Min, Max] | 5.00 [1.00, 5.00] | 5.00 [1.00, 5.00] |
| Missing | 840 (7.1%) | 0 (0%) |
| **Age (months)** |  |  |
| Mean (SD) | 119 (7.50) | 119 (7.47) |
| Median [Min, Max] | 119 [107, 133] | 119 [107, 133] |
| Missing | 1 (0.0%) | 0 (0%) |
| **Pubertal Development** |  |  |
| Pre | 5834 (49.2%) | 3355 (51.3%) |
| Early | 2707 (22.8%) | 1541 (23.6%) |
| Mid/Late | 2853 (24.0%) | 1642 (25.1%) |
| Missing | 474 (4.0%) | 0 (0%) |
| **BMI z-score** |  |  |
| Mean (SD) | 0.407 (1.16) | 0.372 (1.14) |
| Median [Min, Max] | 0.413 [-5.92, 3.07] | 0.367 [-5.92, 3.07] |
| Missing | 77 (0.6%) | 0 (0%) |
| **<sup>a</sup> Race/Ethnicity** |  |  |
| <sup>b</sup> Asian/Other | 1499 (12.6%) | 789 (12.1%) |
| Black | 1784 (15.0%) | 798 (12.2%) |
| Hispanic | 2410 (20.3%) | 1316 (20.1%) |
| White | 6173 (52.0%) | 3635 (55.6%) |
| Missing | 2 (0.0%) | 0 (0%) |
| **<sup>a</sup> Parent Education** |  |  |
| < HS Diploma | 593 (5.0%) | 255 (3.9%) |
| HS Diploma/GED | 1132 (9.5%) | 530 (8.1%) |
| Some College | 3074 (25.9%) | 1589 (24.3%) |
| Bachelor Degree | 3013 (25.4%) | 1713 (26.2%) |
| Post Graduate Degree | 4042 (34.1%) | 2451 (37.5%) |
| Missing/Refused | 14 (0.1%) | 0 (0%) |
| <sup>a</sup> Scanner/Headcoil |  |  |
| Philips Achieva 32-channel | 982 (8.3%) | 591 (9.0%) |
| GE Discovery 32-channel | 2975 (25.1%) | 1324 (20.3%) |
| Philips Ingenia 32-channel | 542 (4.6%) | 236 (3.6%) |
| Siemens Prisma 32-channel | 2634 (22.2%) | 1625 (24.9%) |
| Siemens Prisma 64-channel | 448 (3.8%) | 275 (4.2%) |
| Siemens Prismafit 32-channel | 2915 (24.6%) | 1711 (26.2%) |
| Siemens Prismafit 64-channel | 1372 (11.6%) | 776 (11.9%) |

<sup>a</sup> Difference between research sample and ABCD Study sample is statistically significant at the $p < .05$ level.
<sup>b</sup> The “Other” race/ethnicity category includes participants who were parent-identified as Asian Indian, Chinese, Filipino/a, Japanese, Korean, Vietnamese, Other Asian, American Indian/Native American, Alaska Native, Native Hawaiian, Guamanian, Samoan, Other Pacific Islander, or Other Race.

### Sex

Parent-reported sex is collected as part of the ABCD Study. However, sex assignment at birth is not always an accurate reflection of biological traits like chromosomal sex or gonadal phenotype (19). To more accurately measure the biological influence of sex, we used the frequency ratio of X and Y alleles to define sex according to the presence of a Y chromosome (18) and excluded twelve children whose assigned sex and genetic sex did not match (Figure 1).

### Felt-gender

Participants completed a gender questionnaire at the one-year follow-up visit (17,20). The questionnaire included two questions to measure felt-gender: how much the participant feels like a boy and how much they feel like a girl, on a 5-point Likert scale (“Not at all,” “A little,” “Somewhat,” “Mostly,” “Totally”). Questions were reverse-coded for the gender that was culturally incongruent with the participant’s sex assigned at birth and averaged to create a “felt-gender” score representing the level of congruence between feelings and sex/gender assigned at birth (Supplemental Table 1). A score of 5 indicates feelings congruent with sex assigned at birth, and lower scores reflect higher incongruence.

### Neuroimaging data

The ABCD Study administers a harmonized scanner protocol across 21 data collection sites using a Siemens, GE, or Philips 3T MRI scanner. For a detailed explanation of the imaging protocols, see Casey, et al. (15,16). To reduce motion distortion, real-time, prospective motion correction was implemented. T1w and T2w images were inspected by trained technicians and underwent centralized quality control to identify artifacts or irregularities (16). To improve the accuracy of our deep learning segmentations, we excluded subjects who received a score > 0 for image artifacts or > 1 for motion, inhomogeneity, pial overestimation, or white matter underestimation during the ABCD quality control process (Supplemental Table 2).

Image preprocessing was performed at the Stevens Institute of Neuroimaging and Informatics using the Human Connectome Project minimal preprocessing pipeline to perform regional parcellation and segmentation according to Desikan-Killiany atlas in FreeSurfer 7.1.1, with each white matter region labeled according to the nearest cortical label (21,22). T1w and T2w images were then used to create an enhanced PVS contrast (EPC) image (22). Raters reviewed a single axial slice from each subject to identify EPC images of poor quality that could hinder PVS segmentation and identified a total of 312 additional subjects to be excluded. The EPC images and FreeSurfer parcellations were then used as inputs for our deep learning based Weakly Supervised Perivascular Spaces Segmentation (WPSS) method proposed by Lan et al. (23). WPSS leverages the Frangi filter (24) to enhance and guide the segmentation process, with a particular focus on identifying the salient regions that correspond to the white matter PVS structures. Detailed information about the WPSS procedure can be found in the original publication (23) and in the Supplemental Methods.

After creating PVS masks, we calculated the regional PVS count (number of PVS), volume, and volume fraction for each participant. Given that a high proportion (i.e., > 5%) of subjects had no MR-visible PVS in 12 of the 68 regions from the Desikan-Killiany atlas (Supplemental Table 2), we chose to combine these regions into six larger summary regions of interest (ROIs): centrum semiovale (CSO), the cingulate, and the frontal, temporal, parietal, and occipital lobes. All subsequent statistical analyses were performed on these six ROIs. To account for allometric scaling of volume, descriptive statistics of PVS volume are presented as volume fraction (e.g., regional PVS volume divided by regional white matter volume). However, since scaling of regional metrics is not strictly proportional (25), we adjusted for regional white matter volume as a fixed effect when modeling the relationship between sex, gender, and PVS volume.

### Analysis

All analyses were conducted in R (Supplemental Methods). A model building approach was performed using linear mixed-effects modeling and pairwise ANOVA comparisons of our models to determine the best model of PVS volume and count in each region based on which model explained the most variance in the region of interest (Supplemental Figure 3). Our initial model (M0/) was comprised of pertinent independent variables, including regional volume (in voxels), age (in months, rounded to the nearest whole month), pubertal development status (parent-report), maximum parental education, race/ethnicity (parent-report), BMI z-score, and MRI scanner type as fixed effects and data collection site was included as a random effect (e.g., the nesting of subjects within sites). To capture variation introduced by differences in MRI equipment and software, we created a categorical variable that combined scanner model and the number of head coil channels used (Supplemental Figure 2). Next, we added sex to the initial model to create the sex model (M1), added felt-gender to the initial model to create the felt-gender model (M2), added both sex and felt-gender to create the sex + gender model (M3), and added sex, felt-gender, and their interaction to the initial model to create the interaction model (M4) (Supplemental Table 3).

Since the M3 model contained both sex and felt-gender in addition to all initial model variables, we used Cohen’s f^2^ partial effects sizes to evaluate the relative contribution of each demographic variable to the overall variance in PVS volume and count. To contextualize the practical significance of the sex differences in PVS structure, we further performed a *post hoc* analysis to explore the role of variance on our findings (Supplemental Methods). Lastly, we used the first three principal components from the NIH Toolbox assessment to explore the relationship between PVS and cognition. The first principal component encapsulates general cognitive ability, the second represents executive function, and the third captures learning and memory (26). To test the association of PVS structure on each component of cognition, we ran separate models for volume and count in each of our six ROIs for a total of 12 models for each principal component. The models were created by adding the PVS measure to the sex + gender (M3) model.

## Results

The distribution of mean regional PVS count and volume fraction for each of the 68 Desikan-Killiany atlas regions are depicted in Figure 2, whereas Figure 3 depicts the PVS count and volume fraction for the 6 primary ROIs used in analyses (Supplemental Table 5). The Spearman correlation between count and volume fraction ROIs are shown in Supplemental Figure 3.

**Figure 2.**
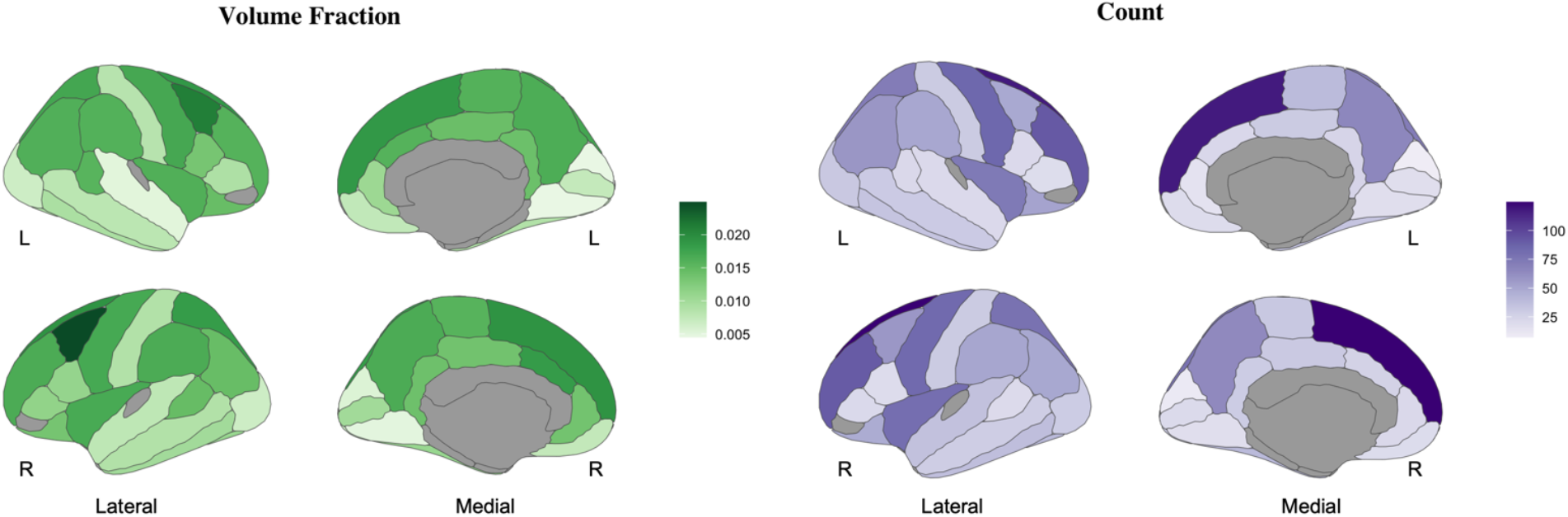
Distribution of PVS volume fraction and count for each Desikan-Killiany atlas region in early adolescents, ages 9-11 years old. Regions where >5% of adolescents had no MR-visible PVS are grayed out.

**Figure 3.**
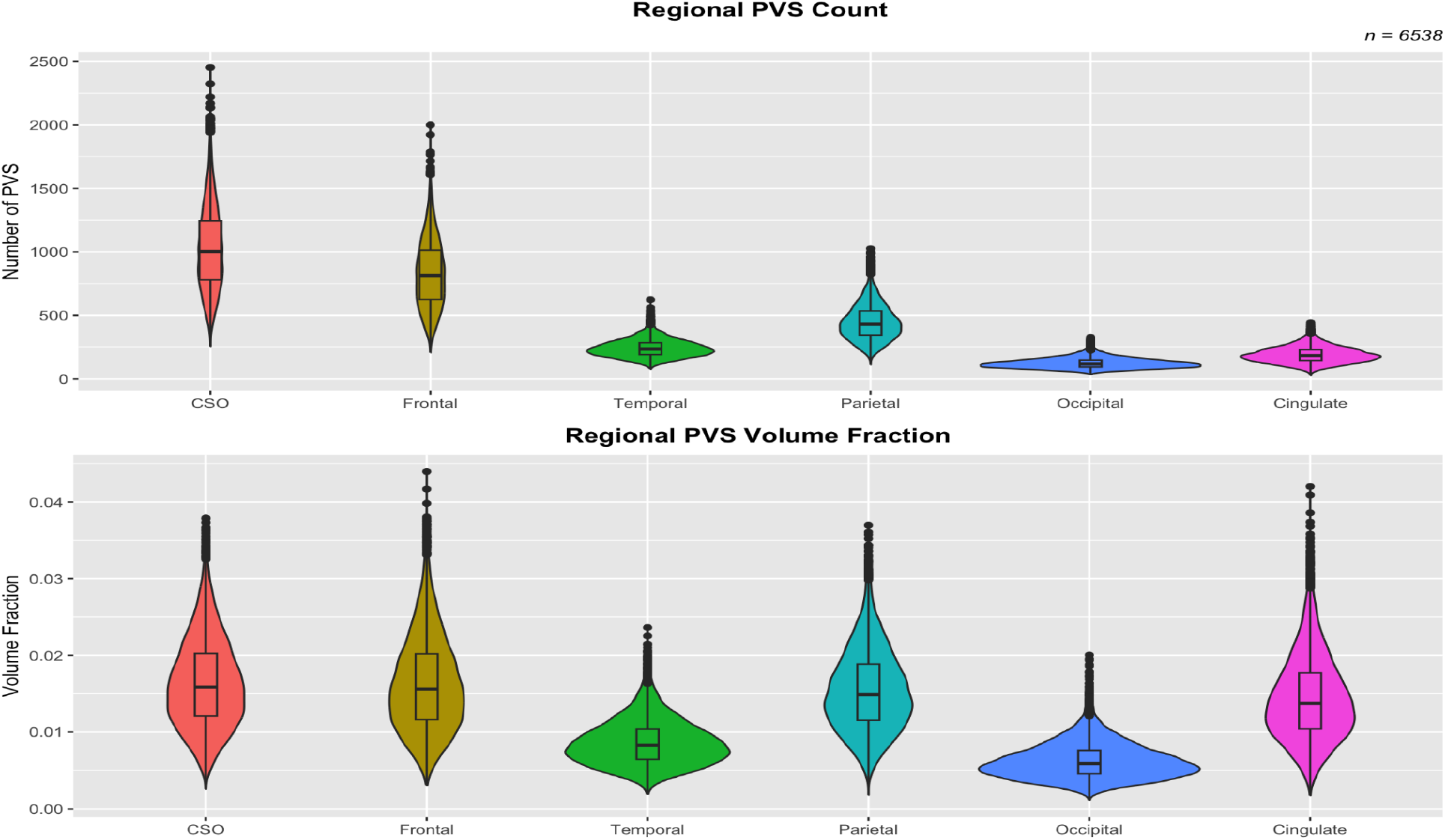
Distribution of PVS volume fraction and count for each of the 6 summary ROIs in early adolescents, ages 9-11 years-old. Abbreviations: CSO = centrum semiovale; PVS = perivascular space.

Pairwise ANOVA comparisons between models determined that the sex model (M1) was the best-fitting model of PVS volume in all ROIs and for PVS count in the temporal lobe. The sex + gender model (M3) was the best fitting model of PVS count in the CSO, frontal lobe, parietal lobe, occipital lobe, and cingulate. The conditional *r*^2^ of the models ranged from 0.342 - 0.477. Model estimates and goodness-of-fit statistics for the PVS volume and count models are provided in Supplemental Tables 6 and 7, respectively. To better understand the relative contributions of demographic factors to PVS structure, we calculated Cohen’s f^2^ partial effect sizes for sex, felt-gender, age, pubertal development, parent education, and BMI z-score (Supplemental Table 8). Of these, only BMI z-score effects met the threshold for a small effect size (Cohen’s f^2^ ≧ 0.02; (27)) (see Figure 5). *Post hoc* findings assessing within- and between-sex variance as well as contextualizing sex differences across the entire sample distribution can be found in the Supplemental Results.

**Figure 5.**
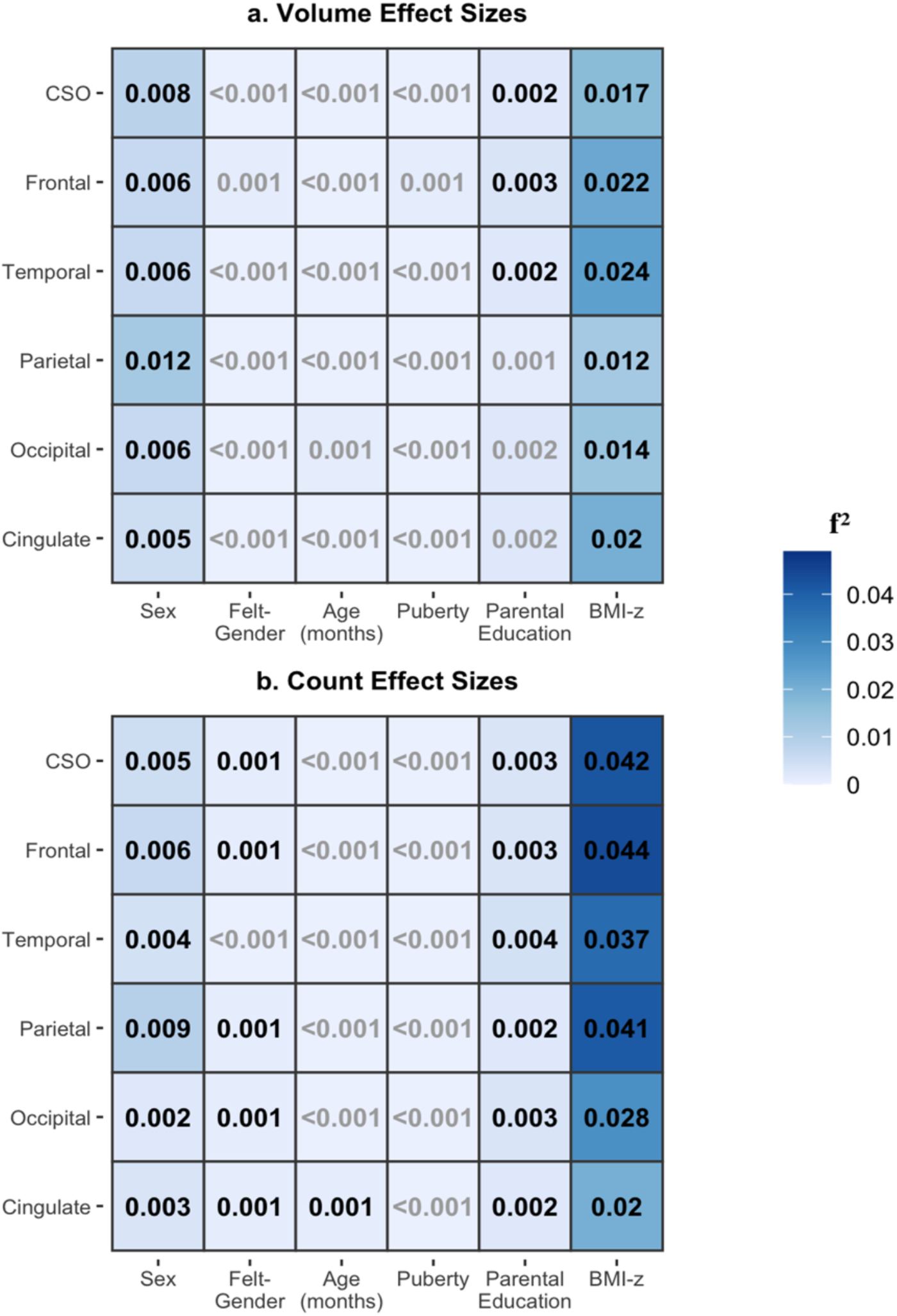
Heat maps of partial effect sizes (Cohen’s partial f^2^) in the sex + gender model (M3) for PVS volume (a) and count (b) in early adolescents, ages 9-11 years-old. Black text indicates that the beta coefficient was significant at the p < 0.05 level. (27). Abbreviations: CSO = Centrum Semiovale

Modeling of the relationship between PVS and cognition revealed that PVS volume was negatively associated with executive function in the cingulate, and with learning/memory in all six ROIs. Similarly, PVS count was negatively associated with cognitive ability in the cingulate, with executive function in the CSO, frontal, temporal, and parietal lobes, and with learning/memory in all six ROIs (Table 4). Although significant relationships were detected between PVS structure and all three principal components of cognition, the partial effect sizes for PVS count and volume were minimal (maximum *f* ^2^ = 0.004). On the other hand, our models were able to account for ∼30% of the overall variance in general cognitive ability (median marginal *r* ^2^ = 0.301; Table 5).

**Table 4.** For each model of the association between PVS structure and the first three principal components of the NIH Toolbox assessment, we provide the beta coefficients, FDR-corrected p-values, and partial effect sizes for regional PVS, as well as the conditional r^2^ of the model. Significant results are highlighted in gray.

| ROI |  | PC1<br>(General Cognitive Ability) |  |  |  | PC2<br>(Executive Function) |  |  |  | PC3<br>(Learning/Memory) |  |  |  |
| --- | --- | --- | --- | --- | --- | --- | --- | --- | --- | --- | --- | --- | --- |
| | | $\beta$ | p-value | partial $f^2$ | $r^2$ | $\beta$ | p-value | partial $f^2$ | $r^2$ | $\beta$ | p-value | partial $f^2$ | $r^2$ |
| Volume | CSO | 0.029 | 0.472 | 1.7e-4 | 0.302 | -0.042 | 0.229 | 2.8e-4 | 0.085 | -0.111 | **2.6e-4** | 0.002 | 0.124 |
|  | Frontal | 0.012 | 0.791 | 3.2e-5 | 0.302 | -0.052 | 0.169 | 4.5e-4 | 0.084 | -0.103 | **4.6e-4** | 0.002 | 0.124 |
|  | Temporal | -0.043 | 0.253 | 4.3e-4 | 0.303 | -0.045 | 0.182 | 3.8e-4 | 0.084 | -0.106 | **2.6e-4** | 0.002 | 0.124 |
|  | Parietal | 0.063 | 0.080 | 8.8e-4 | 0.297 | -0.016 | 0.647 | 4.7e-4 | 0.085 | -0.103 | **3.8e-4** | 0.002 | 0.124 |
|  | Occipital | 0.029 | 0.424 | 2.3e-4 | 0.291 | -2.9e-5 | 0.999 | 1.8e-10 | 0.086 | -0.097 | **2.6e-4** | 0.002 | 0.124 |
|  | Cingulate | -0.059 | 0.080 | 8.1e-4 | 0.299 | -0.074 | **0.034** | 9.8e-4 | 0.085 | -0.090 | **0.001** | 0.002 | 0.124 |
| Count | CSO | -0.023 | 0.709 | 6.4e-5 | 0.302 | -0.143 | **0.003** | 0.002 | 0.086 | -0.158 | **1.2e-4** | 0.003 | 0.125 |
|  | Frontal | -0.042 | 0.424 | 2.2e-4 | 0.302 | -0.144 | **0.003** | 0.002 | 0.086 | -0.147 | **2.6e-4** | 0.003 | 0.125 |
|  | Temporal | -0.099 | 0.065 | 0.001 | 0.304 | -0.112 | **0.032** | 0.001 | 0.085 | -0.196 | **1.4e-5** | 0.004 | 0.126 |
|  | Parietal | 0.007 | 0.917 | 6.6e-6 | 0.297 | -0.119 | **0.013** | 0.001 | 0.086 | -0.161 | **9.9e-5** | 0.003 | 0.125 |
|  | Occipital | -0.002 | 0.949 | 6.7e-7 | 0.291 | -0.057 | 0.182 | 3.6e-4 | 0.086 | -0.132 | **2.6e-4** | 0.002 | 0.124 |
|  | Cingulate | -0.106 | **0.024** | 0.002 | 0.300 | -0.074 | 0.113 | 6.0e-4 | 0.084 | -0.109 | **0.002** | 0.002 | 0.124 |

## Discussion

These findings demonstrate a nuanced relationship between PVS, sex, and felt-gender in 9-11 year-old adolescents. As the first study to report a relationship between gender and PVS, these results also underscore the importance of disentangling the physiological and sociocultural factors in neurovascular health. Notably, the best-fitting model differed between PVS volume and count, suggesting that these variables may represent different physiological aspects of neurovascular structure. For example, count may be more closely related to angiogenesis (28), while volume may be more related to glymphatic function (29). Although sex was related to both count and volume in all ROIs, its effect size was minimal.

Our results also highlight a notable gap in understanding PVS development. Most PVS studies use large ROIs - such as the centrum semiovale, basal ganglia, or total white matter - which provide a large area for counting PVS. When examining PVS in the more fine-grained Desikan-Killiany atlas regions, we found that greater than 5% of participants had no MR-visible PVS in 12/68 regions - particularly within the temporal lobe; however, it is impossible to verify in vivo whether this reflects a true lack of PVS or limitations in image resolution. A prior study with a larger age range (8-21 years) successfully quantified PVS in the Desikan-Killiany atlas ROIs - including in the regions where we observed few MR-visible PVS, but only a few of their participants were as young as our sample (4). Furthermore, their study plotted significant PVS growth during adolescence, with the most change occurring in the temporal lobe (4), suggesting that our findings may be indicative of ongoing regional PVS development.

Consistent with prior research in adults and adolescents (4,6–8), we found that males - on average - have more and larger PVS than females. Nonetheless, these sex differences were minimal in size (Cohen’s f^2^ < 0.02). The physiological basis of this sex difference is likely complex, since both androgen and estrogen receptors are expressed on most cells within the neurovascular unit - including endothelial cells, smooth muscle cells, and astrocytes - and help regulate homeostatic processes such as vasodilation and vasoconstriction, blood-brain barrier permeability, and neuroinflammation, (30,31).

Very few human neuroimaging studies to date have employed a non-dichotomous measure of gender in addition to measuring sex (32,33). Yet, we found that lower congruence between sex and felt-gender was associated with fewer MR-visible PVS in early adolescents. Although the felt-gender effect was minimal, its addition to the linear mixed-effects model did improve model fit, demonstrating that inclusion of gender variables can improve the accuracy of statistical models and account for PVS variance beyond that which can be attributed to sex. Practical interpretation of this result, however, is not straightforward, as the mechanism through which gender and PVS structure is not readily apparent. Future studies should explore the mechanism behind this association by assessing whether differences in PVS are linked to gender-related behavioral differences - like exercise or substance use - or to sociocultural factors like minority stress and allostatic load.

However, our analysis indicated that PVS volume and count both had a larger association with BMI z-score than with sex, felt-gender, age, pubertal development, or parental education. This is in line with research in adults, which similarly found that a higher BMI was associated with a larger PVS volume ratio (8). Like our gender results, more work is needed to ascertain the mechanism through which BMI and PVS are related. For example, childhood obesity is linked to poorer cardiovascular health (34). However, from our results it is unclear whether our BMI results are related to adiposity itself or its downstream health effects.

The importance of understanding sources of variance in PVS structure is reinforced by our finding that both PVS volume and count are associated with multiple aspects of cognition in our sample. Prior research on the cognitive and clinical significance of PVS overwhelmingly focused on adults (10). A recent meta-analysis of PVS and cognition in adults found that higher visual ratings of PVS in the centrum semiovale were associated with worse general intelligence, executive function, language, and memory (12). In contrast to the interpretation of such findings in adults as a potential biomarker of neurodegeneration or age-related cognitive decline (10,12), we found that both PVS count and volume were negatively associated with early adolescent learning and memory scores in all ROIs and higher PVS count was associated with lower executive function in the centrum semiovale, frontal, temporal, and parietal lobes. These broad findings of a relationship between PVS structure and cognition at ages 9-11 suggests that the relationship between PVS, learning, and memory persists across the lifespan and emphasizes the importance of studying normative PVS structure, function and development.

## Limitations

The generalizability of these results is hampered by our sample composition. Despite aiming to employ probability sampling to the extent that it was feasible, the ABCD Study exhibits over-representation of white participants and highly educated parents (35). Our data exclusion and quality control process further exacerbated this bias (see Table 1). Excluding participants for missing data and poor image quality can introduce bias because image quality and exclusion criteria are related to a broad spectrum of behavioral, demographic, and health--related variables (36). More work is needed to improve participation of under-represented socioeconomic, racial, and ethnic groups in publicly available datasets.

As a cross-sectional study, this research captures only a snapshot of early adolescent PVS development. Although our results are in line with other studies of adolescent PVS, no study to date has examined PVS development longitudinally prior to age 12. As such, we cannot ascertain whether PVS absence in subregions of the temporal lobe reflects biological immaturity or technical limitations. For example, the temporal lobe is plagued by signal distortion and dropout (37). Similarly, although effect sizes offer valuable standardized interpretations, context is also important. Negligible effects are relevant in certain contexts, and cumulative or long-term impacts may be more meaningful. In fact, small effect sizes are to be expected in brain research in particular (38).

The use of gender in human health and physiology research is still an emerging field. While we focused on felt-gender due to its non-dichotomous nature and availability in a large, nationwide sample, gender itself is complex and multifaceted. The present study was unable to capture all aspects of gender identity, expression, or interpersonal relationships that might contribute to adolescent brain structure. Furthermore, the gender questionnaire was not administered at the baseline imaging visit and imaging was not collected at the one-year follow-up, so we relied on gender responses collected approximately one year after imaging. Therefore, our study assumes felt-gender is relatively stable over a ∼12-month period between ages of 9-11. Future research should collect multiple dimensions of gender concurrent with MR-imaging to improve precision.

## Conclusions

This research study demonstrates that both sex and felt-gender have significant impacts on PVS structure in early adolescence, albeit with minimal effect sizes. BMI z-score had a stronger relationship with PVS count and volume than sex, felt-gender, age, pubertal development, or parental education. Even after accounting for all of these demographic variables and regional white matter volume, PVS count and volume showed a significant, negative association with learning and memory in all regions examined. PVS count also showed a significant negative relationship with executive function in 4/6 regions. Overall, this study illustrates the importance of studying the development of PVS and the relationship between PVS structure and healthy brain function.

## Supporting information

Supplemental Results

## Notes

https://abcdstudy.org

### Competing Interest Statement

Jeiran Choupan declares her status as an employee of the private company NeuroScope Inc (NY, USA)

### Funding

- National Institutes of Health: U01DA041048, U01DA050989, U01DA051016, U01DA041022, U01DA051018, U01DA051037, U01DA050987, U01DA041174, U01DA041106, U01DA041117, U01DA041028, U01DA041134, U01DA050988, U01DA051039, U01DA041156, U01DA041025, U01DA041120, U01DA051038, U01DA041148, U01DA041093, U01DA041089, U24DA041123, and U24DA041147
- National Institute of Mental Health: RF1MH123223

## References

1. Wardlaw JM, Benveniste H, Nedergaard M, et al. Perivascular spaces in the brain: anatomy, physiology and pathology. Nat Rev Neurol. Nature Publishing Group; 2020;16(3):137–153.

2. Karkoska KA, Gollamudi J, Sawyer RP, Woo D, Hyacinth HI. Quantifying dilated perivascular spaces in children with sickle cell disease. Pediatr Blood Cancer. 2024;71(9):e31150. doi: 10.1002/pbc.31150.

3. Park M-G, Roh J, Ahn S-H, Cho JW, Park K-P, Baik SK. Dilated perivascular spaces and steno-occlusive changes in children and adults with moyamoya disease. BMC Neurol. 2024;24(1):14. doi: 10.1186/s12883-023-03520-z.

4. Lynch KM, Sepehrband F, Toga AW, Choupan J. Brain perivascular space imaging across the human lifespan. Neuroimage. Elsevier; 2023;271:120009.

5. Kim HG, Shin N-Y, Nam Y, et al. MRI-visible Dilated Perivascular Space in the Brain by Age: The Human Connectome Project. Radiology. 2023;306(3):e213254. doi: 10.1148/radiol.213254.

6. Piantino J, Boespflug EL, Schwartz DL, et al. Characterization of MR Imaging– Visible Perivascular Spaces in the White Matter of Healthy Adolescents at 3T. Am J Neuroradiol. 2020;41(11):2139–2145. doi: 10.3174/ajnr.A6789.

7. Yamamoto EA, Koike S, Wong C, et al. Biological sex and BMI influence the longitudinal evolution of adolescent and young adult MRI-visible perivascular spaces. bioRxiv. 2024;2024.08.17.608337. doi: 10.1101/2024.08.17.608337.

8. Barisano G, Sheikh-Bahaei N, Law M, Toga AW, Sepehrband F. Body mass index, time of day and genetics affect perivascular spaces in the white matter. J Cereb Blood Flow Metab Off J Int Soc Cereb Blood Flow Metab. 2021;41(7):1563–1578. doi: 10.1177/0271678X20972856.

9. Zhang C, Chen Q, Wang Y, et al. Risk Factors of Dilated Virchow-Robin Spaces Are Different in Various Brain Regions. Hendrikse J, editor. PLoS ONE. 2014;9(8):e105505. doi: 10.1371/journal.pone.0105505.

10. Francis F, Ballerini L, Wardlaw JM. Perivascular spaces and their associations with risk factors, clinical disorders and neuroimaging features: A systematic review and meta-analysis. Int J Stroke. SAGE Publications; 2019;14(4):359–371. doi: 10.1177/1747493019830321.

11. Streed CG, Caceres BA, Mukherjee M. Preventing cardiovascular disease among sexual and gender minority persons. Heart Br Card Soc. 2021;107(13):1100–1101. doi: 10.1136/heartjnl-2021-319069.

12. Liu L, Tu L, Shen Q, et al. Meta-analysis of the relationship between the number and location of perivascular spaces in the brain and cognitive function. Neurol Sci Off J Ital Neurol Soc Ital Soc Clin Neurophysiol. 2024;45(8):3743–3755. doi: 10.1007/s10072-024-07438-3.

13. Gouveia-Freitas K, Bastos-Leite AJ. Perivascular spaces and brain waste clearance systems: relevance for neurodegenerative and cerebrovascular pathology. Neuroradiology. 2021;63(10):1581–1597. doi: 10.1007/s00234-021-02718-7.

14. Voorter PHM, van Dinther M, Jansen WJ, et al. Blood–Brain Barrier Disruption and Perivascular Spaces in Small Vessel Disease and Neurodegenerative Diseases: A Review on MRI Methods and Insights. J Magn Reson Imaging. 2024;59(2):397–411. doi: 10.1002/jmri.28989.

15. Casey BJ, Cannonier T, Conley MI, et al. The Adolescent Brain Cognitive Development (ABCD) study: Imaging acquisition across 21 sites. Dev Cogn Neurosci. 2018;32:43–54. doi: 10.1016/j.dcn.2018.03.001.

16. Hagler DJ, Hatton SN, Cornejo MD, et al. Image processing and analysis methods for the Adolescent Brain Cognitive Development Study. NeuroImage. 2019;202:116091. doi: 10.1016/j.neuroimage.2019.116091.

17. Potter AS, Dube SL, Barrios LC, et al. Measurement of gender and sexuality in the Adolescent Brain Cognitive Development (ABCD) study. Dev Cogn Neurosci. 2022;53:101057. doi: 10.1016/j.dcn.2022.101057.

18. Torgerson C, Ahmadi H, Choupan J, Fan CC, Blosnich JR, Herting MM. Sex, gender diversity, and brain structure in early adolescence. Hum Brain Mapp. 2024;45(5):e26671. doi: 10.1002/hbm.26671.

19. Joel D. Genetic-gonadal-genitals sex (3G-sex) and the misconception of brain and gender, or, why 3G-males and 3G-females have intersex brain and intersex gender. Biol Sex Differ. 2012;3(1):27. doi: 10.1186/2042-6410-3-27.

20. Potter A, Dube S, Allgaier N, et al. Early adolescent gender diversity and mental health in the Adolescent Brain Cognitive Development (ABCD) study. J Child Psychol Psychiatry. 2021;62(2):171–179. doi: 10.1111/jcpp.13248.

21. Desikan RS, Ségonne F, Fischl B, et al. An automated labeling system for subdividing the human cerebral cortex on MRI scans into gyral based regions of interest. NeuroImage. 2006;31(3):968–980. doi: 10.1016/j.neuroimage.2006.01.021.

22. Glasser MF, Sotiropoulos SN, Wilson JA, et al. The minimal preprocessing pipelines for the Human Connectome Project. NeuroImage. 2013;80:105–124. doi: 10.1016/j.neuroimage.2013.04.127.

23. Lan H, Lynch KM, Custer R, et al. Weakly supervised perivascular spaces segmentation with salient guidance of frangi filter. Magn Reson Med. 2023;89(6):2419–2431. doi: 10.1002/mrm.29593.

24. Frangi AF, Niessen WJ, Vincken KL, Viergever MA. Multiscale Vessel Enhancement Filtering..

25. Sanchis-Segura C, Ibañez-Gual MV, Aguirre N, Cruz-Gómez ÁJ, Forn C. Effects of different intracranial volume correction methods on univariate sex differences in grey matter volume and multivariate sex prediction. Sci Rep. Nature Publishing Group; 2020;10(1):12953. doi: 10.1038/s41598-020-69361-9.

26. Sukumaran K, Botternhorn KL, Schwartz J, et al. Associations between Fine Particulate Matter Components, Their Sources, and Cognitive Outcomes in Children Ages 9–10 Years Old from the United States. Environ Health Perspect. Environmental Health Perspectives; 2024;132(10):107009. doi: 10.1289/EHP14418.

27. Cohen J. A power primer. Psychol Bull. 1992;112(1):155–159. doi: 10.1037//0033-2909.112.1.155.

28. Wälchli T, Bisschop J, Carmeliet P, et al. Shaping the brain vasculature in development and disease in the single-cell era. Nat Rev Neurosci. 2023;24(5):271– 298. doi: 10.1038/s41583-023-00684-y.

29. Yu L, Hu X, Li H, Zhao Y. Perivascular Spaces, Glymphatic System and MR. Front Neurol. 2022;13:844938. doi: 10.3389/fneur.2022.844938.

30. Gonçalves FJ, Abrantes-Soares F, Pouso MR, Lorigo M, Cairrao E. Non-genomic Effect of Estradiol on the Neurovascular Unit and Possible Involvement in the Cerebral Vascular Accident. Mol Neurobiol. 2023;60(4):1964–1985. doi: 10.1007/s12035-022-03178-7.

31. Lucas-Herald AK, Touyz RM. Androgens and Androgen Receptors as Determinants of Vascular Sex Differences Across the Lifespan. Can J Cardiol. 2022;38(12):1854–1864. doi: 10.1016/j.cjca.2022.09.018.

32. Dhamala E, Bassett DS, Yeo BT, Holmes AJ. Functional brain networks are associated with both sex and gender in children. Sci Adv. 2024;10(28):eadn4202. doi: 10.1126/sciadv.adn4202.

33. Rauch JM, Eliot L. Breaking the binary: Gender versus sex analysis in human brain imaging. NeuroImage. 2022;264:119732. doi: 10.1016/j.neuroimage.2022.119732.

34. Friedemann C, Heneghan C, Mahtani K, Thompson M, Perera R, Ward AM. Cardiovascular disease risk in healthy children and its association with body mass index: systematic review and meta-analysis. The BMJ. 2012;345:e4759. doi: 10.1136/bmj.e4759.

35. Osayande N, Marotta J, Aggarwal S, et al. Diversity-aware Population Models: Quantifying Associations between Socio-Spatial Factors and Cognitive Development in the ABCD Cohort. Res Sq. 2024;rs.3.rs-4751673. doi: 10.21203/rs.3.rs-4751673/v1.

36. Peverill M, Russell JD, Keding TJ, et al. Balancing Data Quality and Bias: Investigating Functional Connectivity Exclusions in the Adolescent Brain Cognitive Development^SM^ (ABCD Study) Across Quality Control Pathways. Hum Brain Mapp. 2025;46(1):e70094. doi: 10.1002/hbm.70094.

37. Olman CA, Davachi L, Inati S. Distortion and Signal Loss in Medial Temporal Lobe. García AV, editor. PLoS ONE. 2009;4(12):e8160. doi: 10.1371/journal.pone.0008160.

38. Owens MM, Potter A, Hyatt CS, et al. Recalibrating expectations about effect size: A multi-method survey of effect sizes in the ABCD study. PLOS ONE. Public Library of Science; 2021;16(9):e0257535. doi: 10.1371/journal.pone.0257535.

