## Supplemental Methods for "Sex and gender differences in perivascular space in early adolescence"

### Participants

Data for this study came from the ongoing Adolescent Brain Cognitive Development (ABCD) Study® (Casey et al., 2018; Hagler et al., 2019). Children were excluded from the ABCD Study if they had a lack of English proficiency, severe sensory, neurological, medical or intellectual limitations, or inability to complete an MRI scan (Li et al., 2021). In terms of age, sex, and household size, the ABCD cohort was selected to closely match the distribution of 9-11-year-olds in the American Community Survey (Heeringa & Berglund, 2020). We retrieved the raw T1-weighted and T2-weighted structural MRI files from the publicly available NDA 3.0 release (NDA 3.0 data release 2020; <https://dx.doi.org/10.15154/1520591>) and used tabulated questionnaire data obtained from the NDA 5.0 release (NDA 5.0 data release 2023; <https://nda.nih.gov/study.html?id=2147>).

### Neuroimaging

FreeSurfer segmentation quality metrics range from 0 to 3, with scores of zero indicating no image quality issues (i.e., no susceptibility artifact).To improve the accuracy of our deep learning segmentations, we excluded subjects who received a score greater than 0 for image artifacts or a score greater than 1 for motion, inhomogeneity, pial overestimation, or white matter underestimation in the tabulated FreeSurfer quality control data provided by the ABCD Study.

After preprocessing we combined T1- and T2-weighted images to create an enhanced PVS contrast (EPC) image to improve our ability to detect PVS, as outlined by Sepehrband, et al. (Sepehrband et al., 2019). Raters then reviewed a single axial slice from each subject to identify EPC images with poor image quality that could hinder PVS segmentation (i.e. inhomogeneity, artefact) and identified a total of 323 additional subjects to be excluded. The EPC images and FreeSurfer parcellations were then used as inputs for our Weakly Supervised Perivascular Spaces Segmentation (WPSS) method proposed by Lan et al. (Lan et al., 2023). WPSS leverages the Frangi filter (Frangi et al., 1998) to enhance and guide the segmentation process, with a particular focus on identifying the salient regions that correspond to the PVS structures. WPSS employs weak supervision by incorporating salient features derived from the Frangi filter. These features enable the model to learn from fewer labeled data, thereby reducing the need for extensive manual annotation while still achieving high segmentation accuracy. Initially, a convolutional neural network algorithm U-Net (Ronneberger et al., 2015) is trained to differentiate between perivascular spaces and surrounding tissues. Following the initial segmentation produced by the U-Net, a Conditional Random Field (CRF) as a Recurrent Neural Network (RNN) module (Zheng et al., 2015) is employed to further refine the PVS segmentation by leveraging the salient guidance from the Frangi filter. This dual-step approach allows the model to achieve precise segmentations, even with limited annotated data. The WPSS model used for this ABCD study was trained on the HCP dataset (Bookheimer et al., 2019; Somerville et al., 2018; Van Essen et al., 2013), using 400 enhanced PVS contrast (EPC) images generated by the pipeline proposed by Sepehrband et al. (Sepehrband et al., 2019). Detailed information about the training data and the quality control procedure can be found in the original publication (Lan et al., 2023). A total of 136 participants failed to complete the data processing pipeline due to an error in pre-processing, FreeSurfer segmentation, or PVS segmentation and were excluded.

### Mixed-Effects Modeling

We applied a model building approach using linear mixed-effects modeling via the lme4 and lmerTest packages in R (Bates et al., 2015; Kuznetsova et al., 2017; R Core Team, 2019a) to determine the best model of PVS volume and count in each region. To account for other sources of neuroanatomical variance beyond sex and gender, our null model (M0̷) was comprised on pertinent independent variables, including regional volume, age, pubertal development status, maximum parental education, race/ethnicity, BMI z-score, and MRI scanner type as fixed effects and data collection site was included as a random effect (e.g., the nesting of subjects within sites). Age was recorded in months and rounded to the nearest whole month. Pubertal development was based on the parent-report version of the Pubertal Development Scale (PDS) and categorized as prepuberty, early puberty, mid puberty, late puberty, and post-puberty (Cheng et al., 2021; Petersen et al., 1988). Given the low numbers of participants in the late or post-pubertal categories, we combined the mid, late, and post-puberty groups into a single category (mid/late puberty). Due to systemic social injustice, ecological factors that contribute to neurodevelopment may be correlated with sociocultural variables like race, ethnicity, and socioeconomic status (Nketia et al., 2021; Werchan & Amso, 2017). Therefore, we chose to include a measure of combined race and ethnicity (White, Hispanic, Black, Other) and the maximum level of parent education (less than high school diploma, high school diploma or GED, some college, bachelor’s degree, or postgraduate degree) in the null model (M0̷). To capture variation introduced by differences in MRI equipment and software, we created a categorical variable that combined scanner model and the number of head coil channels used (Siemens Prisma 32-channel, Siemens Prisma 64-channel, Siemens Prisma fit 32-channel, Siemens Prisma fit 64-channel, Philips Ingenia 32-channel, GE Discovery 32-channel, Philips Achieva 32-channel) ([Supplemental Figure 2](https://docs.google.com/document/d/10UkmEyzdgf2BzGPbJBunp8W2hf_1hTlPkdime4QoNOc/edit?tab=t.0#heading=h.xxsbttlnu9xs)).

To verify that the assumptions of linear mixed-effects models were not violated we visually examined the residuals of each model. After running all 5 models for each region, we used pairwise ANOVA comparisons to determine which model best explained the variance in the region of interest. If no model fit the outcome variance significantly better than the null model (M0̷), then we considered the null model the best-fitting model. For the interaction model (M4) to be the best-fitting model, we required it to have a p-value < 0.05 in the ANOVA comparisons to all 4 other models (M0̷, M1, M2, and M3). The criteria for the sex + gender model (M3) to be labeled the best-fitting model was that it had a p-value < 0.05 in ANOVA comparisons with the M0̷, M1, and M2 models, and the interaction model was not the best model. If none of the above criteria were met and only one of the remaining models (M1 or M2) significantly improved fit compared to the null model (M0̷), then the model that performed better than the null was considered the best model for explaining outcome variance. If both the sex model (M1) and the gender model (M2) had a p-value < 0.05 when compared to the null model (M0̷), then the model with the lowest Akaike Information Criterion (AIC) was chosen as the best-fitting model for the data.

To explore the relative contribution of each of our demographic variables to the overall variance in PVS structure, we compared the Cohen’s f^2^ partial effect sizes from the sex + gender (M3) model, since it contained all of our independent variables of interest other than the sex-by-felt-gender interaction - which was not a significant contributor to PVS volume or count in any region.

### *Post Hoc* Analyses

To contextualize the findings of average differences in PVS structure between male and female adolescents, we further tested for sex differences in variance and examined the differences in PVS across the entire sample distribution. As previously published (Torgerson et al., 2024), we used a series of tests to assess inhomogeneity of variance, similarity, and effect size using the unadjusted as well as adjusted (i.e. residuals from M0) PVS counts and volume (Torgerson et al., 2024). For assessing similarity, we used the overlap coefficient (OVL_2_) using the bayestestR package (Markowski et al., 2019). The Fligner-Kileen test was run using the stats package (R Core Team, 2019b) to assess regional inhomogeneity of variance (e.g., significant differences in variance between male and female adolescents). Cliff’s delta test, from the effsize package (Torchiano, 2020), was performed to examine the effect size for sex differences across the entire data distribution (Cliff, 2014). From the dominance matrix calculated for the Cliff’s delta test, we also obtained the probability that a randomly selected male would have a greater PVS volume or count than a randomly selected female (also known as the common language effect size or CLES)(Mastrich & Hernandez, 2021). FDR correction was used to adjust the p-values from the mixed-effects models and the Fligner-Killeen test.
