## Supplemental Results for "Sex and gender differences in perivascular space in early adolescence"

### ***Post Hoc*** Analyses

We found a minimum of 73.6% overlap between male and female distributions for all unadjusted PVS measures, and a minimum of 89.7% overlap after adjustment for other sources of variance (i.e. M0). The Fligner-Killeen test yielded significant sex differences in all regions examined, with greater male than female variance. Cliff’s δ found small and medium effect sizes across the sample distribution for PVS volume and count before adjustment. However, after adjustment effect sizes for sex were minimal in magnitude, suggesting that other variables - such as BMI - are contributing to the observed sex differences in the unadjusted data. Adjustment also reduced the CLES. Prior to adjustment, this probability ranged from 62.1% - 67.7%, but after adjustment, this probability was close to chance (range: 51.4% - 54.3%).

| PVS Volume | | Centrum  Semiovale | Frontal | Temporal | Parietal | Occipital | Cingulate |
| --- | --- | --- | --- | --- | --- | --- | --- |
| Unadjusted | Overlap | 0.760 | 0.779 | 0.744 | 0.745 | 0.803 | 0.807 |
|  | Fligner-Killeen | 82.2 | 74.1 | 113.4 | 92.9 | 79.0 | 43.5 |
|  | Fligner-Killeen  p-value | **1.2E-19** | **7.5E-18** | **1.8E-26** | **5.6E-22** | **6.1E-19** | **4.2E-11** |
|  | Cliff's δ | 0.344 | 0.319 | 0.358 | 0.357 | 0.273 | 0.275 |
|  | Cliff's  magnitude | medium | small | medium | medium | small | small |
|  | CLES | 0.672 | 0.659 | 0.679 | 0.678 | 0.636 | 0.637 |
| Adjusted | Overlap | 0.908 | 0.912 | 0.909 | 0.892 | 0.921 | 0.934 |
|  | Fligner-Killeen | 72.4 | 57.3 | 88.4 | 99.4 | 68.4 | 40.1 |
|  | Fligner-Killeen  p-value | **1.7E-17** | **3.8E-14** | **5.4E-21** | **2.0E-23** | **1.3E-16** | **2.5E-10** |
|  | Cliff's δ | 0.069 | 0.065 | 0.049 | 0.086 | 0.050 | 0.051 |
|  | Cliff's  magnitude | negligible | negligible | negligible | negligible | negligible | negligible |
|  | CLES | 0.535 | 0.533 | 0.525 | 0.543 | 0.525 | 0.526 |
| PVS Count | | Centrum  Semiovale | Frontal | Temporal | Parietal | Occipital | Cingulate |
| Unadjusted | Overlap | 0.803 | 0.812 | 0.734 | 0.777 | 0.824 | 0.816 |
|  | Fligner-Killeen | 26.5 | 25.1 | 63.1 | 41.7 | 18.0 | 21.1 |
|  | Fligner-Killeen  p-value | **2.7E-07** | **5.3E-07** | **2.0E-15** | **1.1E-10** | **2.2E-05** | **4.3E-06** |
|  | Cliff's δ | 0.300 | 0.290 | 0.378 | 0.326 | 0.252 | 0.257 |
|  | Cliff's  magnitude | small | small | medium | small | small | small |
|  | CLES | 0.650 | 0.644 | 0.687 | 0.662 | 0.622 | 0.626 |
| Adjusted | Overlap | 0.925 | 0.929 | 0.930 | 0.924 | 0.952 | 0.949 |
|  | Fligner-Killeen | 21.8 | 20.4 | 42.6 | 41.1 | 23.4 | 14.3 |
|  | Fligner-Killeen  p-value | **3.0E-06** | **6.2E-06** | **6.6E-11** | **1.4E-10** | **1.3E-06** | **1.6E-04** |
|  | Cliff's δ | 0.066 | 0.068 | 0.052 | 0.079 | 0.033 | 0.047 |
|  | Cliff's  magnitude | negligible | negligible | negligible | negligible | negligible | negligible |
|  | CLES | 0.533 | 0.534 | 0.526 | 0.539 | 0.516 | 0.524 |

#### ***Table 4.*** *Results for unadjusted and adjusted (residualized) PVS volume and count in early adolescents ages 9-11 years-old. Statistical tests include: Overlap Coefficient Fligner-Killeen test for inhomogeneity of variance, Cliff’s δ, and the CLES (the probability that a randomly selected male would have a greater PVS volume or count than a randomly selected female).*
