## Supplemental Figures for "Sex and gender differences in perivascular space in early adolescence"

### **Supplemental Figure 1.** A violin plot showing the residuals from the null model for the centrum semiovale (CSO) by scanner and head coil type.

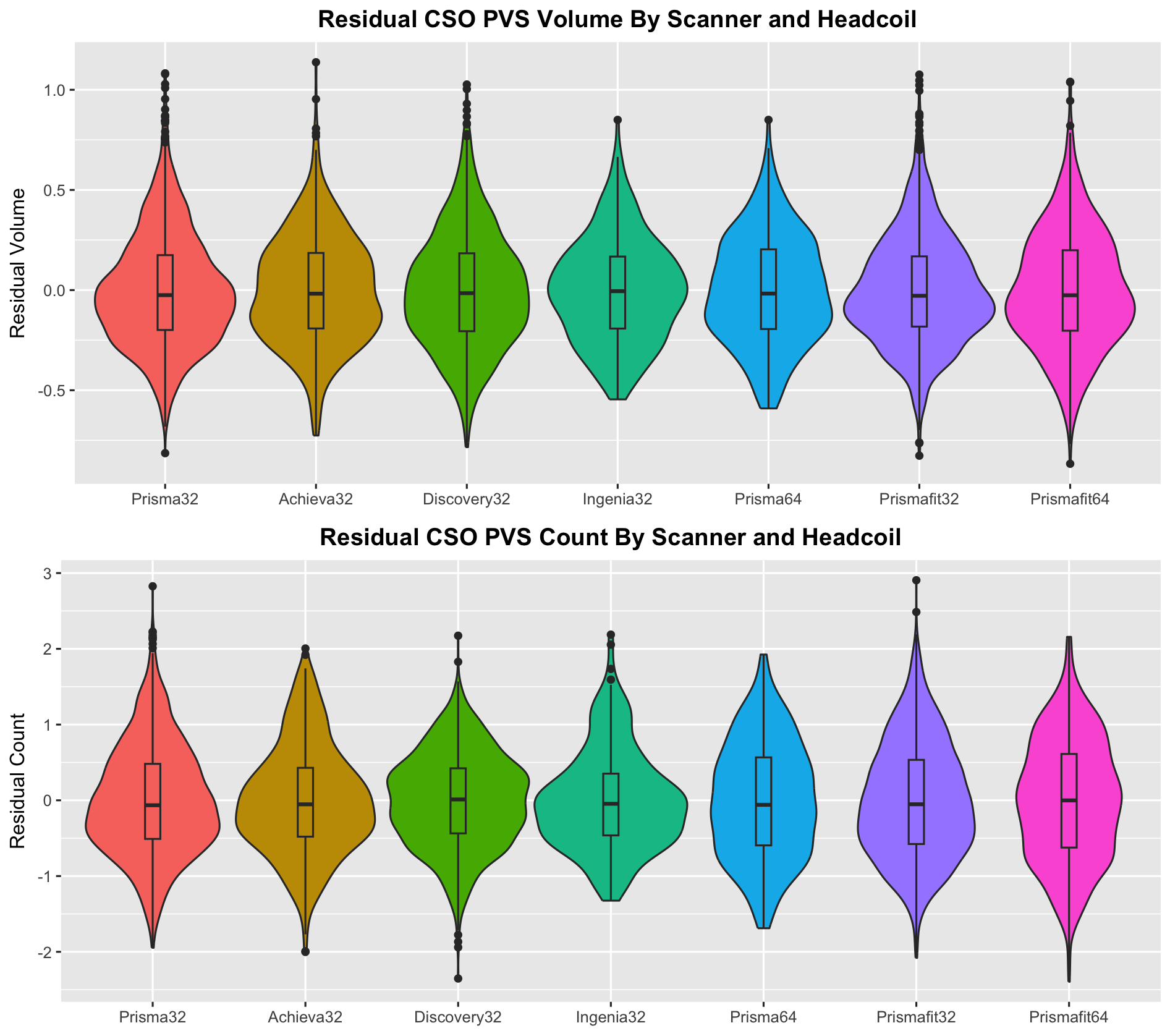

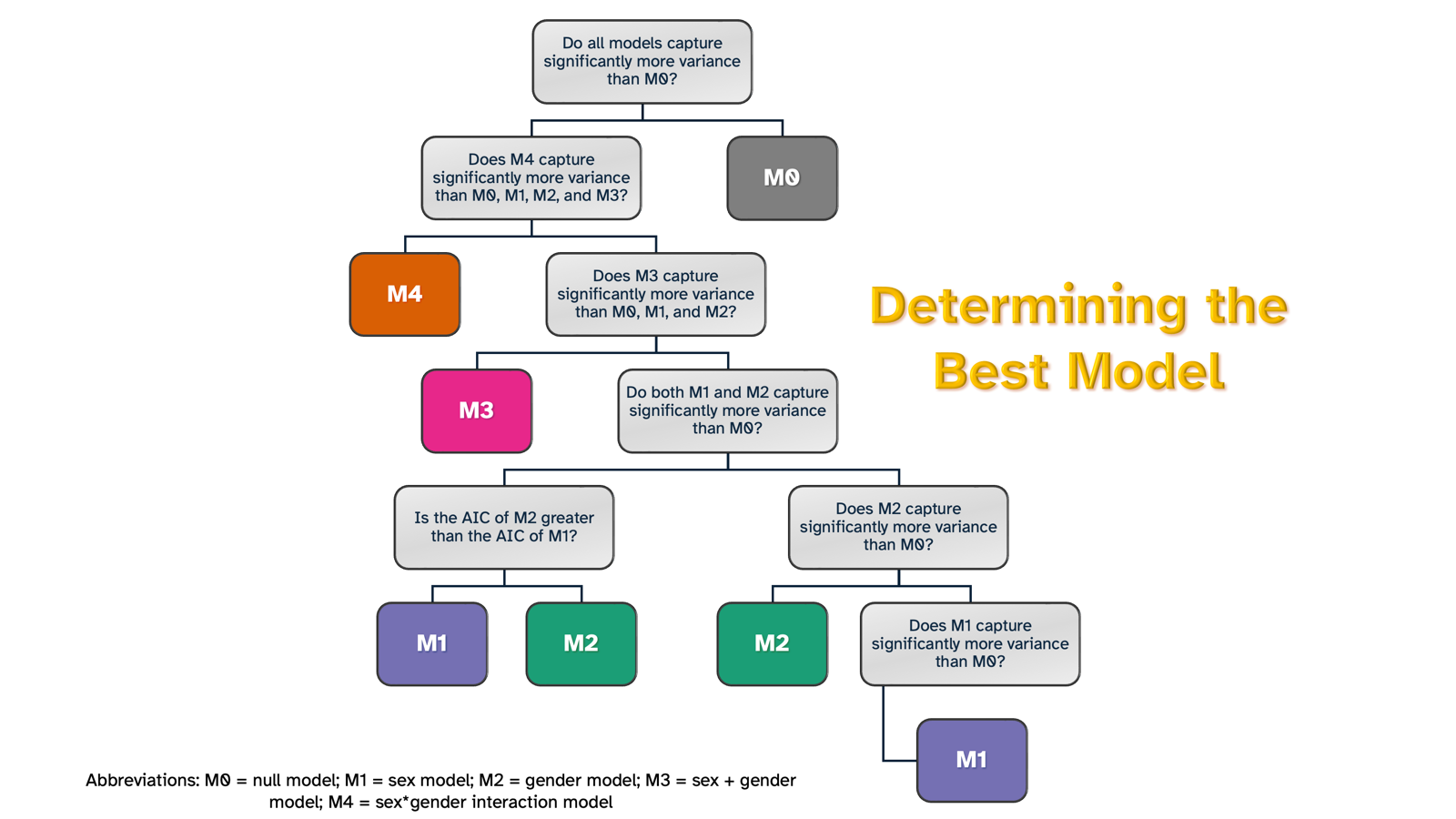

### **Supplemental Figure 2.** A decision tree demonstrating the criteria used to determine the best-fitting model after performing pairwise ANOVA comparisons of all models for each region.

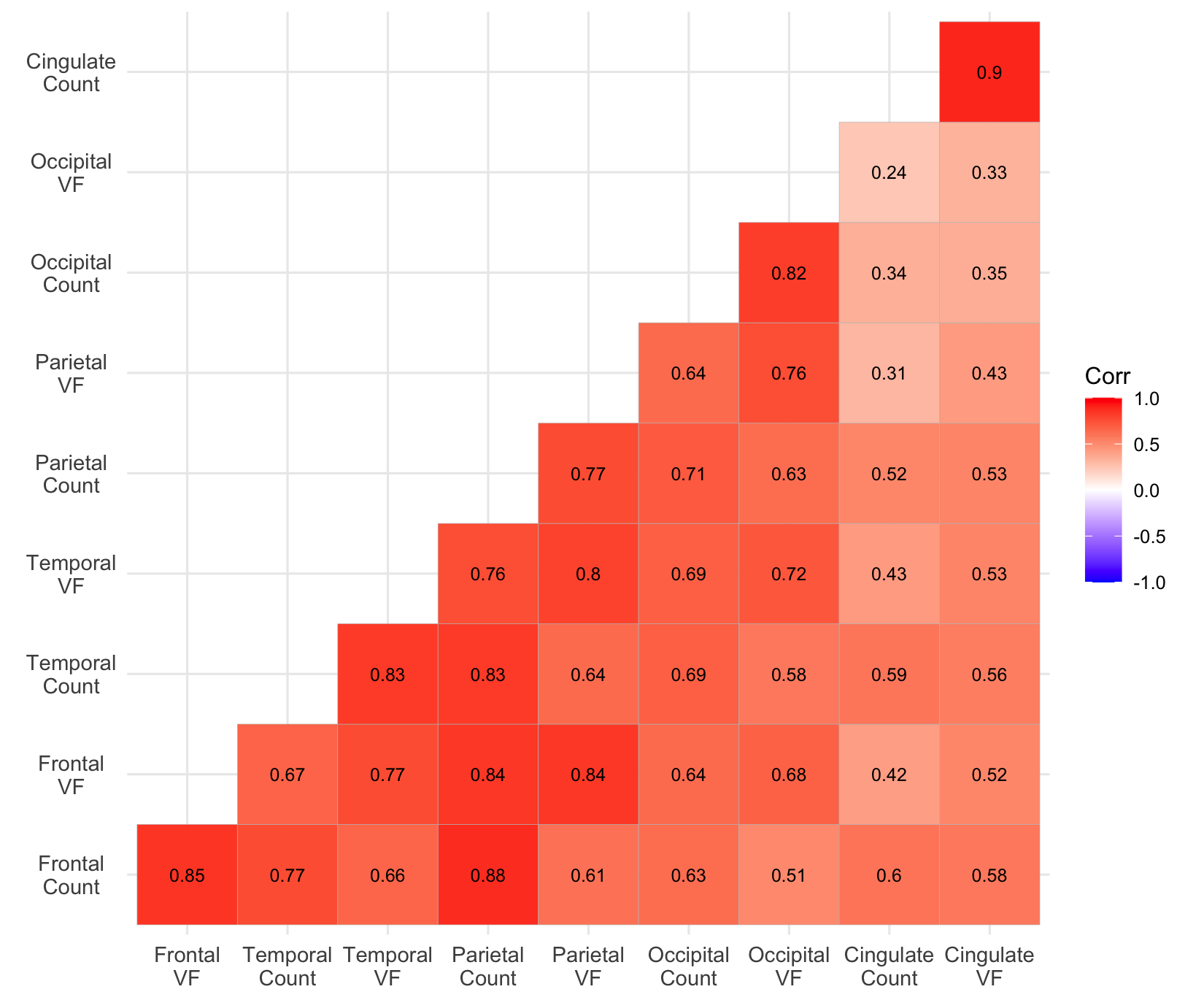

### **Supplemental Figure 3.** A correlation plot showing the Spearman’s correlation between each PVS count and volume fraction ROI; abbreviations: VF = volume fraction.
